# Replication protein A prevents unregulated priming and Rad51 loading on single-stranded DNA in nuclear extracts of *Xenopus* eggs

**DOI:** 10.64898/2026.06.09.731035

**Authors:** Taisei Miyata, Naoki Tani, Yoshitaka Kawasoe, Kei-ichiro Ishiguro, Tatsuro S. Takahashi

## Abstract

In eukaryotes, single-stranded DNA (ssDNA) generated during DNA replication, recombination, and repair is rapidly bound and protected by the major single-stranded DNA-binding protein replication protein A (RPA). RPA not only stabilizes ssDNA but also acts as a central platform that coordinates diverse DNA transactions. Exhaustion of RPA due to unregulated ssDNA production leads to replication fork breakage and replication catastrophe, underscoring its critical role in genome stability. However, the direct consequences of RPA limitation remain incompletely understood. Using *Xenopus* egg extracts, we show that excess ssDNA induces spontaneous priming, a reaction that is otherwise prevented in a physiological nuclear environment. We provide evidence that priming suppression is mediated by stoichiometric binding of RPA to ssDNA. Analysis of the ssDNA-binding proteome reveals that RPA promotes the association of ATR checkpoint factors, Polα–primase, and the RFWD3 ubiquitin ligase with ssDNA. In contrast, RPA depletion induces the recruitment of Rad51, Rad51 paralogs, and Fbh1, a DNA helicase that interacts with both RPA and Rad51 and promotes fork breakage under replication stress. Collectively, our findings suggest that RPA contributes to genome stability by protecting ssDNA from inappropriate DNA synthesis and unscheduled recruitment of recombination and fork-processing factors.

## Introduction

Replication protein A (RPA), also known as replication factor A (RF-A), is the primary single-stranded DNA (ssDNA)-binding protein in eukaryotes. In most eukaryotic cells, the major form of RPA is a heterotrimer composed of RPA1, RPA2, and RPA3 (Wold 1997; Fanning et al. 2006; Caldwell and Spies 2020). The RPA complex contains six oligonucleotide/oligosaccharide-binding (OB) folds, designated DNA-binding domains (DBDs) A through F; DBD-A, -B, -C, and - F reside in RPA1, whereas DBD-D and -E are located in RPA2 and RPA3, respectively. DBD-A through D primarily mediate ssDNA binding of the RPA complex, enabling flexible DNA-binding modes that vary in both length and affinity, ranging from 8 to 30 nucleotides (Fanning et al. 2006; Caldwell and Spies 2020; Ahmad et al. 2021). Because of its high abundance and overall subnanomolar ssDNA-binding affinity, ssDNA is rapidly and extensively coated by RPA upon exposure in eukaryotic cells.

ssDNA is an essential and unavoidable intermediate in diverse DNA transactions, including DNA replication, recombination, and repair. Reflecting its importance, RPA not only protects ssDNA from nucleolytic degradation and secondary structure formation but also coordinates these processes by serving as a versatile protein-protein interaction platform (Marechal and Zou 2015; Bhat and Cortez 2018; Dueva and Iliakis 2020). RPA was identified as an essential factor for SV40 *in vitro* replication (Wobbe et al. 1987; Fairman and Stillman 1988; Wold and Kelly 1988), and studies using this system have provided key insights into its functions (Waga and Stillman 1998). RPA stimulates the viral helicase large T antigen to promote origin unwinding (Fairman and Stillman 1988; Wold and Kelly 1988), and subsequent studies showed that it is also required for origin unwinding during cellular DNA replication (Walter and Newport 2000; Yeeles et al. 2015), although this function does not appear to depend on specific interactions with the CMG replicative helicase (Pike et al. 2023). Studies in the SV40 system further showed that RPA stimulates priming and DNA synthesis by the DNA polymerase α-primase (Polα-primase) complex and DNA polymerase δ–proliferating cell nuclear antigen (PCNA)–replication factor C (Waga and Stillman 1998). At saturating concentrations, however, RPA inhibits Polα-primase (Collins and Kelly 1991; Tsurimoto and Stillman 1991; Taylor and Yeeles 2018), and this suppression is alleviated through interactions between Polα-primase and replicative DNA helicases in both SV40 and cellular systems (Weisshart et al. 2000; Huang et al. 2010; Jones et al. 2023). Beyond replication, RPA also plays central roles in homologous recombination and DNA repair (Dueva and Iliakis 2020). In homologous recombination, DNA double-strand breaks (DSBs) are resected from 5′ to 3′ to generate 3′ ssDNA tails required for strand exchange. RPA promotes DSB end resection by stimulating the nuclease Dna2, protects exposed ssDNA from homology-based annealing and secondary structure formation, and subsequently, the RPA-coated ssDNA serves as the substrate for Rad51 nucleoprotein filament formation (Kowalczykowski 2015). RPA is also an indispensable component of nucleotide excision repair and DNA mismatch repair (Jiricny 2013; Kuper and Kisker 2026).

Moreover, RPA-coated ssDNA functions as a critical signaling platform for the ATR checkpoint pathway, which governs both replication stress and the DNA damage response (Marechal and Zou 2015; Dueva and Iliakis 2020). The central component of this pathway, the ATR-ATRIP complex, binds RPA-coated ssDNA (Zou and Elledge 2003; Bomgarden et al. 2004). The ATR kinase is activated by the checkpoint mediators TOPBP1 and ETAA1 and the sensor complex RAD9–HUS1–RAD1 (9–1–1), all of which interact with RPA (Zou and Elledge 2003; Kumagai et al. 2006; Marechal and Zou 2015; Bass et al. 2016; Feng et al. 2016; Haahr et al. 2016; Lee et al. 2016). Interestingly, ATR prevents excess ssDNA exposure by suppressing dormant origin firing under replication stress (Toledo et al. 2013). If this function is compromised, the nuclear pool of RPA is exhausted upon the accumulation of ssDNA by unregulated origin firing, leading to replication fork breakage and a replication catastrophe. These findings indicate that stoichiometric protection of ssDNA by RPA is critical for controlling cellular DNA metabolism, and that when ssDNA is generated in excess, it becomes vulnerable to aberrant protein binding and inappropriate DNA transactions. While the terminal phenotype of RPA exhaustion is characterized by global DNA breakage, the immediate biochemical transitions on ssDNA that occur when RPA becomes limiting remain uncertain.

In this study, we used nuclear extracts from *Xenopus* eggs, a physiological *in vitro* system that recapitulates DNA replication, recombination, and repair, to examine the molecular events caused by insufficient RPA supply.

## Results

### Excess ssDNA induces spontaneous priming in the nucleoplasmic extract from *Xenopus* eggs

A demembranated interphase extract of unfertilized *Xenopus* eggs, called high-speed supernatant (HSS), supports origin licensing (Mcm2–7 loading onto dsDNA), *de novo* priming on ssDNA, and second-strand DNA synthesis, but not replication of dsDNA (Mechali and Harland 1982; Riedel et al. 1982). Initiation of DNA replication in this system depends on nuclear formation, which concentrates replication factors within the nucleus. A highly concentrated extract of nuclear proteins, called nucleoplasmic extracts (NPE), recapitulates a replication-competent nuclear environment and thereby supports efficient firing of licensed origins and subsequent dsDNA replication (Walter et al. 1998). Notably, although HSS supports spontaneous priming on ssDNA, NPE does not (Walter and Newport 2000; Stokes et al. 2002). The absence of spontaneous priming in NPE suggests that DNA synthesis is tightly regulated at the priming step in a functional nuclear environment.

Nuclear proteins are approximately 10- to 30-fold enriched in NPE compared with HSS (Walter et al. 1998). We therefore hypothesized that priming inhibition may be mediated by enrichment of a priming inhibitor in NPE. If such an inhibitor functions in a dose-dependent manner, excess ssDNA might titrate it out, thereby permitting spontaneous priming in NPE. To test this idea, we incubated circular phagemid ssDNA in NPE at increasing concentrations (Fig. 1). DNA loading was adjusted such that each lane contained the same total amount of DNA, irrespective of the ssDNA concentration during incubation. DNA synthesis was monitored by supplying NPE with Cy5-dUTP. At 10 ng/μL ssDNA, we faithfully recapitulated previous findings (Walter and Newport 2000; Stokes et al. 2002); a single pre-annealed primer allowed nearly complete conversion of ssDNA into dsDNA, whereas no DNA synthesis was detected in the absence of a pre-annealed primer, indicating that the priming step is blocked in NPE (Fig. 1, lanes 1–4). Strikingly, at ssDNA concentrations above 20 ng/μL, we observed robust DNA synthesis in the absence of pre-annealed primers. The observed incorporation was sensitive to aphidicolin, suggesting that B-family polymerases, most likely DNA polymerase δ, mediate strand extension. These results strongly suggest that a dose-dependent inhibitor suppresses spontaneous priming on ssDNA in NPE.

**Figure 1.**
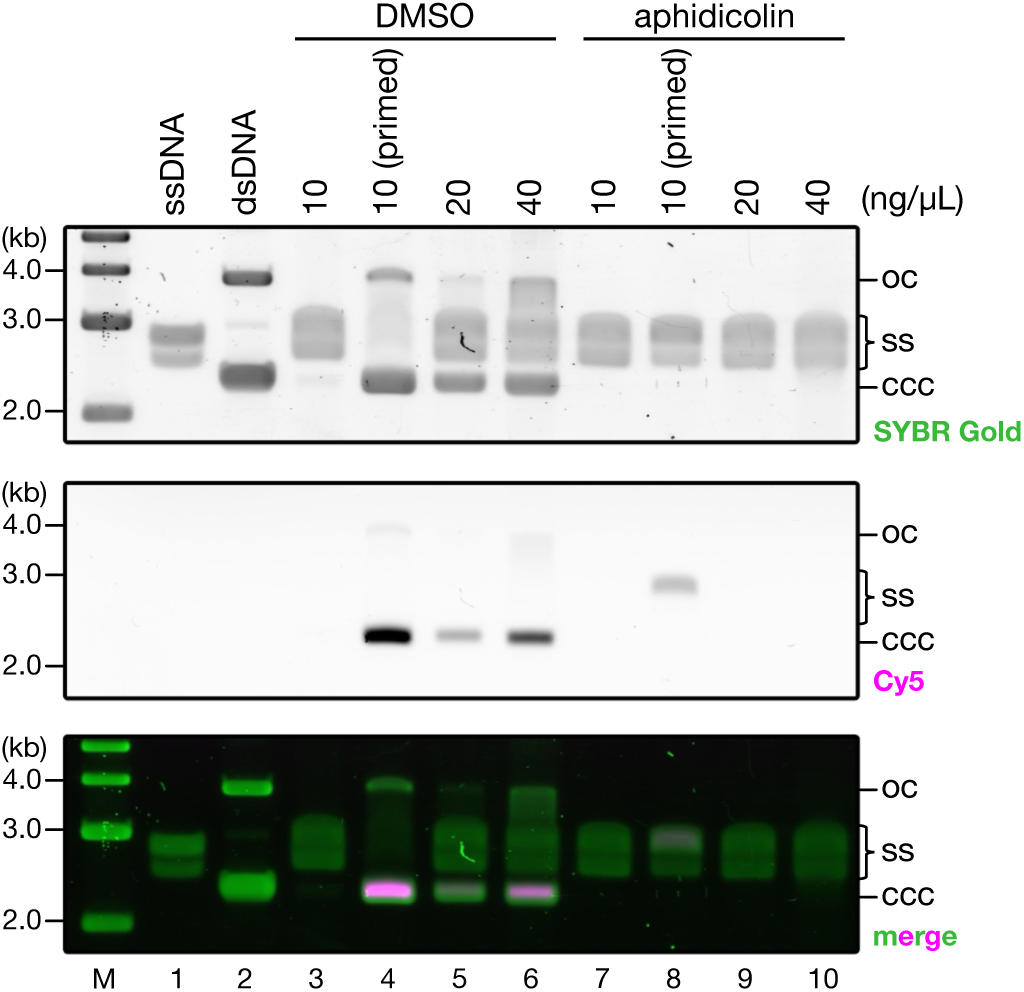
Excess ssDNA induces spontaneous priming in NPE. Circular ssDNA derived from pBluescript KS(−) was incubated in NPE supplemented with 25 μM Cy5-dUTP at the indicated concentrations for 2 h in the absence (lanes 3, 5–7, 9, and 10) or presence (lanes 4 and 8) of a pre-annealed primer. Where indicated, aphidicolin (50 ng/μL; lanes 7–10) or vehicle (lanes 3–6) was added. DNA samples were purified and separated by 0.8% agarose gel electrophoresis. *M*, DNA size marker; *oc*, open circular DNA; *ss*, single-stranded DNA; *ccc*, covalently closed circular DNA.

### RPA is responsible for priming suppression

Given its high abundance and strong affinity for ssDNA, RPA is a strong candidate for the inhibitory activity that suppresses spontaneous priming in NPE. Consistent with this idea, primer synthesis on ssDNA by Polα–primase is inhibited by saturating concentrations of RPA *in vitro* (Collins and Kelly 1991; Tsurimoto and Stillman 1991; Weisshart et al. 2000; Huang et al. 2010; Taylor and Yeeles 2018). We therefore tested whether RPA is responsible for the suppression of spontaneous priming in NPE.

To this end, we co-expressed *Xenopus* Rpa1–3 in insect cells and purified the Rpa1–3 heterotrimer in a nearly stoichiometric form (Figs. 2A and B). The recombinant heterotrimer was used both to generate antibodies against Rpa1–3 and as a standard for quantitative immunoblotting. Using this approach, we estimated the endogenous concentration of Rpa1 in NPE to be approximately 5 μM (Fig. 2C). Because RPA occupies approximately 8–30 nucleotides of ssDNA depending on its binding mode, this concentration of RPA is predicted to stoichiometrically protect ∼40–150 μM nucleotides of ssDNA, corresponding to approximately 13–50 ng/μL ssDNA. Thus, ssDNA concentrations in the range of 20–40 ng/μL are expected to approach or exceed the capacity of NPE to protect ssDNA.

**Figure 2.**
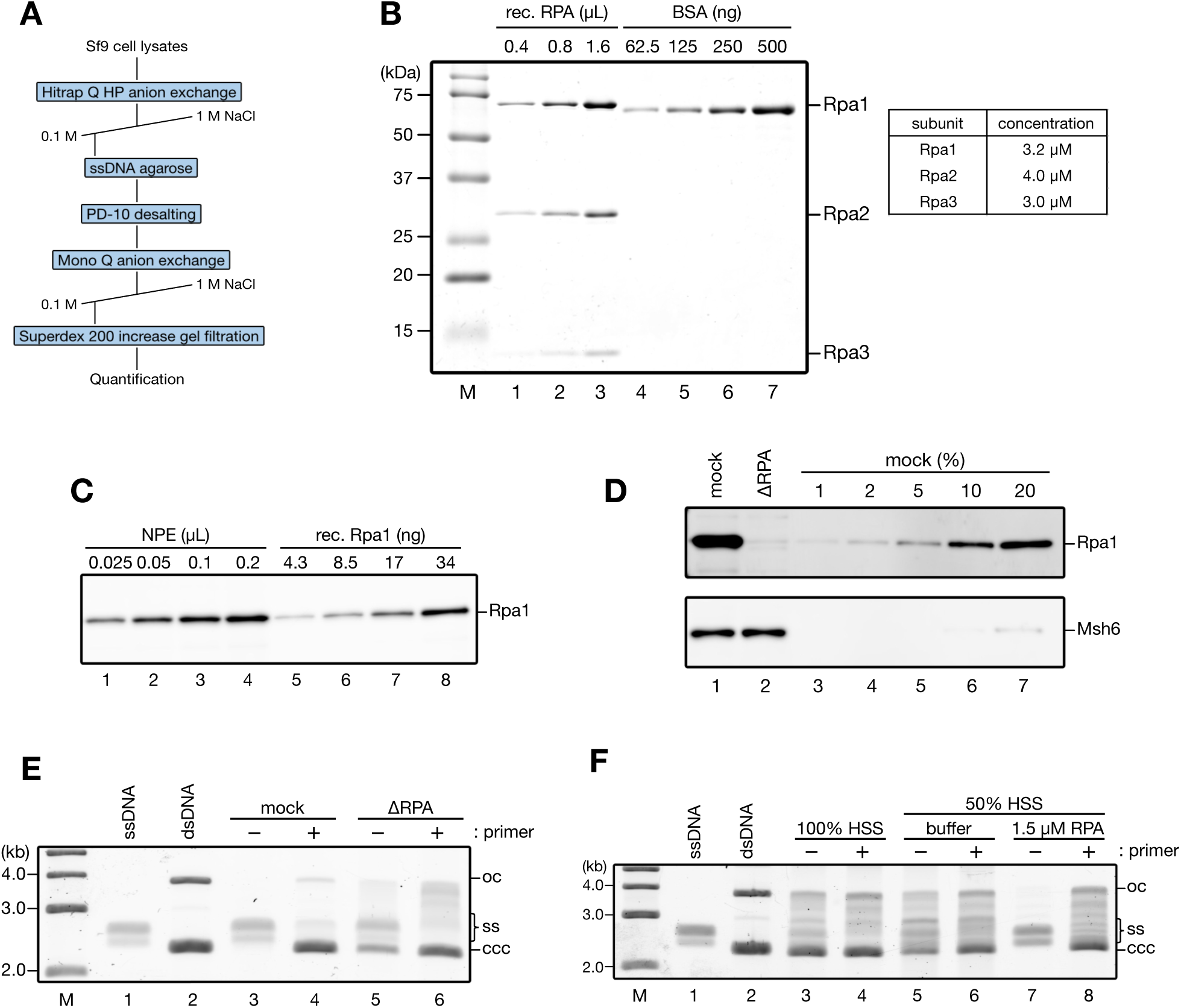
RPA is responsible for priming suppression. (A) Schematic representation of the purification procedure for *Xenopus* RPA. (B) Purified recombinant RPA heterotrimer was separated by SDS-PAGE, stained with Coomassie Brilliant Blue R-250, and quantified using bovine serum albumin (BSA) as a standard. Estimated concentrations of individual subunits are shown in the table, from which the concentration of the RPA heterotrimer was estimated to be approximately 3.0 μM. *M*, protein size marker. (C) The indicated amounts of NPE and recombinant RPA were separated by SDS-PAGE and probed with RPA antibody. Comparison with recombinant RPA standards yielded an estimated endogenous Rpa1 concentration of approximately 5 μM in NPE. (D) 0.2 μL each of mock-treated NPE (lane 1), RPA-depleted NPE (lane 2), and a serial dilution series of mock-treated NPE (lanes 3–7) were analyzed by immunoblotting with the indicated antibodies. Msh6 served as a loading control. (E) Circular ssDNA was incubated in the NPE characterized in (D) at 10 ng/μL for 2 h in the absence (lanes 3 and 5) or presence (lanes 4 and 6) of a pre-annealed primer. DNA samples were purified and separated by 0.8% agarose gel electrophoresis. *M*, DNA size marker; *oc*, open circular DNA; *ss*, single-stranded DNA; *ccc*, covalently closed circular DNA. (F) Circular ssDNA was incubated at 10 ng/μL in undiluted (100%) or 50%-diluted HSS supplemented with buffer (lanes 5 and 6) or 1.5 μM recombinant RPA (lanes 7 and 8) for 2 h in the absence (lanes 3, 5, and 7) or presence (lanes 4, 6, and 8) of a pre-annealed primer. DNA samples were purified and separated by 0.8% agarose gel electrophoresis.

Guided by this estimate, we next depleted endogenous RPA from NPE using RPA antibodies (Fig. 2D). Immunodepletion was >98% efficient for Rpa1, resulting in a ∼50-fold reduction in the capacity of NPE to protect ssDNA. In mock-treated extracts, spontaneous priming was suppressed at 10 ng/μL ssDNA (Fig. 2E). In contrast, DNA synthesis was observed even in the absence of a pre-annealed primer in RPA-depleted extracts, which are predicted to retain a maximal ssDNA protection capacity of only ∼0.2–0.8 ng/μL. These findings strongly suggest that RPA is responsible for suppressing spontaneous priming on ssDNA in NPE.

As an independent approach, we supplied recombinant RPA to HSS to test whether it could suppress spontaneous priming normally observed in this extract (see Fig. 2F, lane 3). To maximize the concentration of recombinant RPA, HSS was mixed with recombinant RPA at a 1:1 ratio. HSS diluted to 50% with the buffer, which serves as a control, supported DNA synthesis on 10 ng/μL ssDNA in the absence of pre-annealed primers (Fig. 2F). Because HSS contains approximately 20-fold less RPA than NPE, the ssDNA protection capacity of 50%-diluted HSS is predicted to be approximately 0.3–1.3 ng/μL. Strikingly, addition of 1.5 μM recombinant RPA, which corresponds to an additional ssDNA protection capacity of approximately 4–15 ng/μL, completely suppressed DNA synthesis. Together, these results demonstrate that RPA is the factor responsible for priming suppression in *Xenopus* egg extracts.

### ssDNA-binding proteomes in the presence and absence of RPA

The cellular capacity to protect ssDNA is inherently limited by RPA abundance. This principle underlies the RPA exhaustion model, which predicts that excess ssDNA leads to a functional shortage of RPA, triggering replication stress that ultimately results in DNA breakage (Toledo et al. 2013; Toledo et al. 2017). Our results indicate that one consequence of RPA exhaustion in *Xenopus* egg extracts is unregulated priming on ssDNA. However, unregulated priming alone is unlikely to fully account for the DNA damage associated with RPA exhaustion.

To explore additional processes that may contribute to genome instability when RPA becomes limiting, we analyzed how RPA depletion affects the ssDNA-binding proteome in NPE. A biotinylated 90-mer ssDNA oligonucleotide was immobilized on streptavidin-coated beads and incubated in mock-treated or RPA-depleted NPE. Bead-bound proteins were then recovered and analyzed by mass spectrometry (Figs. 3A–C and Table S1). Silver staining confirmed ssDNA-dependent recovery of multiple proteins in mock-treated extracts, whereas RPA depletion markedly reduced association of RPA subunits (Fig. 3C). Consistent with previous observations (You et al. 2002; Lee et al. 2003; Recolin et al. 2012; Lyu et al. 2019), mass spectrometry revealed a substantial reduction of Atr, Atrip, and Etaa1, ATR pathway components whose ssDNA association is RPA-dependent, from the ssDNA-binding proteome following RPA depletion (Fig. 3D). Rfwd3, another known RPA interactor and RPA-dependent ssDNA-binding protein (Elia et al. 2015; Feeney et al. 2017; Inano et al. 2017), was also significantly underrepresented in RPA-depleted extracts. Recruitment of Pola1 and Prim2, the DNA polymerase α–primase subunits, was similarly reduced, suggesting that loading of Pol α–primase is at least partially dependent on RPA. Together, these results demonstrate how RPA shapes the ssDNA-binding proteome in *Xenopus* egg extracts.

**Figure 3.**
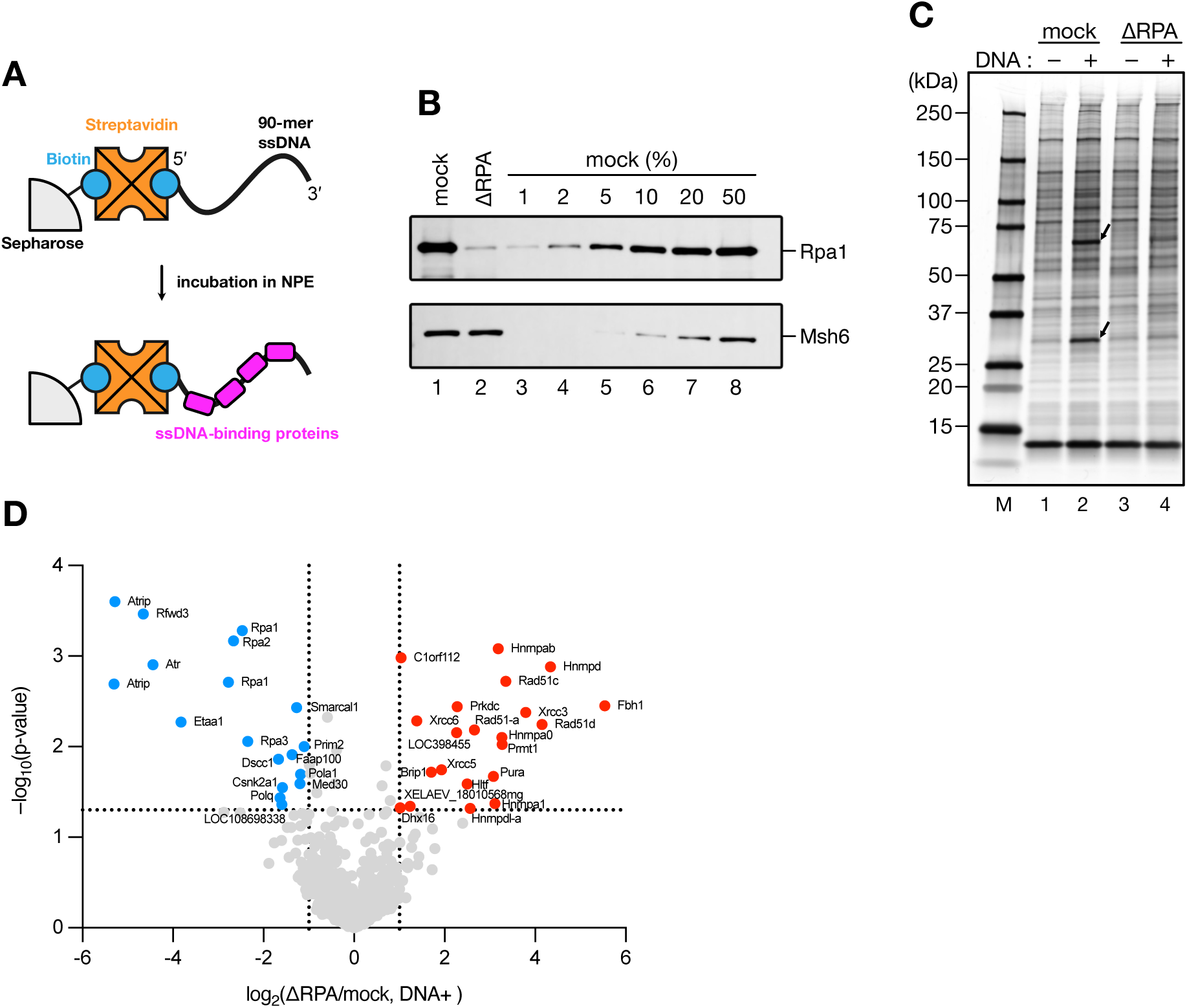
ssDNA-binding proteomes in the presence and absence of RPA. (A) Schematic representation of the ssDNA pulldown assay. A 5′-biotinylated 90-mer ssDNA was first mixed with streptavidin (SA), and the resulting DNA–SA complex was subsequently immobilized on biotinylated Sepharose beads. The DNA-coated beads were incubated in NPE at 22°C for 10 min. (B) 0.2 μL each of mock-treated NPE (lane 1), RPA-depleted NPE (lane 2), and a serial dilution series of mock-treated NPE (lanes 3–8) were analyzed by immunoblotting with the indicated antibodies. Msh6 served as a loading control. (C) The ssDNA pulldown assay was performed using the NPE characterized in (B) in the absence (lanes 1 and 3) or presence (lanes 2 and 4) of immobilized ssDNA. Bead-bound proteins were separated by SDS-PAGE and visualized by silver staining. Arrows indicate Rpa1 (67 kDa) and Rpa2 (29 kDa). (D) Volcano plot of log_2_ fold changes between RPA-depleted and mock-treated conditions in the presence of ssDNA versus −log_10_ p-values derived from three independent experiments. Dashed lines indicate p = 0.05 and log_2_ fold change = ±1.

### RPA suppresses recruitment of homologous recombination factors

Our proteomic analysis also identified multiple ssDNA-associated factors that became enriched upon RPA depletion. Notably, many of these proteins are linked to homologous recombination. In particular, ssDNA association of the Rad51 recombinase was markedly increased in RPA-depleted extracts, accompanied by enrichment of Rad51 paralogs and Rad51 regulators, including Rad51C, Rad51D, Xrcc3, and Fbh1 (Morishita et al. 2005; Osman et al. 2005; Ito et al. 2024). These findings suggest that loss of RPA promotes unscheduled activation of the Rad51 pathway on ssDNA.

To validate these observations, we analyzed ssDNA-bound proteins by immunoblotting. Consistent with efficient depletion, RPA association with ssDNA was strongly reduced in RPA-depleted extracts (Fig. 4). Under these conditions, Rad51 binding to ssDNA increased significantly: Rad51 was not detectable on ssDNA in mock-treated extracts, whereas robust Rad51 association was observed following RPA depletion. Together, these results suggest that RPA protects ssDNA by preventing unscheduled Rad51 loading.

**Figure 4.**
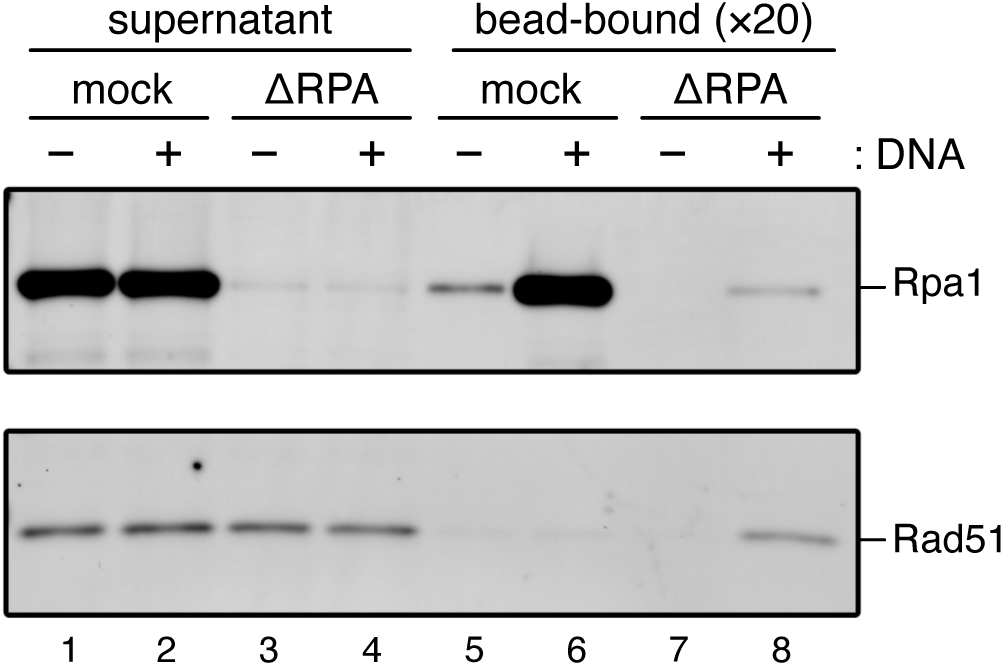
RPA depletion promotes Rad51 loading onto ssDNA. The ssDNA pulldown assay described in Figure 3A was performed in the absence (lanes 1, 3, 5, and 7) or presence (lanes 2, 4, 6, and 8) of immobilized ssDNA. Supernatants (lanes 1–4) and bead-bound proteins (lanes 5–8) were separated by SDS-PAGE and probed with the indicated antibodies.

## Discussion

In this study, we used *Xenopus* egg extracts to examine the direct consequences of RPA limitation. Our results identify RPA as a dose-dependent inhibitor of spontaneous priming on ssDNA. Thus, depletion of RPA from NPE activated priming and DNA synthesis, whereas addition of recombinant RPA to HSS suppressed spontaneous priming. By comparing proteomes on ssDNA in the presence and absence of RPA, we further identified factors whose ssDNA loading is either promoted or suppressed by RPA. Collectively, these findings provide mechanistic insight into how RPA protects ssDNA in a functional nuclear environment and reinforce its role as a key regulator of priming and Rad51 loading.

The ability of RPA to inhibit primer synthesis has been documented in the SV40 and yeast *in vitro* replication systems (Collins and Kelly 1991; Tsurimoto and Stillman 1991; Weisshart et al. 2000; Taylor and Yeeles 2018), where interactions between Polα–Primase and DNA helicases alleviate the inhibitory effect (Weisshart et al. 2000; Huang et al. 2010; Jones et al. 2023). Our findings extend these principles to a physiological nuclear context and suggest that, while RPA promotes the recruitment of Polα–primase, it also acts as a potent inhibitor of its priming activity, thereby restricting priming to conditions in which RPA is locally displaced, for example, by DNA helicases. Such regulation is likely important during homologous recombination, as inappropriate spontaneous priming would counteract end resection by converting ssDNA back to dsDNA.

It is well known that priming is readily observed on ssDNA in HSS, a crude extract of interphase egg cytoplasm and nucleoplasm (Mechali and Harland 1982; Riedel et al. 1982). Since the amphibian cell cycle involves open mitosis during which the nucleoplasm disperses into the cytoplasm, HSS reflects an interphase-like environment prior to functional nuclear assembly upon mitotic exit. The permissiveness of HSS for spontaneous priming suggests that ssDNA may be vulnerable to unscheduled priming during open mitosis.

Our data also highlight the dual role of RPA as both a protector of ssDNA and a platform for regulated protein recruitment. Consistent with previous studies, loading of Atr–Atrip and Etaa1 is dependent on RPA, reinforcing the established model in which RPA-coated ssDNA activates the ATR checkpoint (Marechal and Zou 2015; Acevedo et al. 2016; Bass et al. 2016; Feng et al. 2016; Haahr et al. 2016). Rfwd3, an RPA-interacting factor (Elia et al. 2015; Feeney et al. 2017; Inano et al. 2017), similarly depends on RPA for its association with ssDNA. In contrast, Rad51 loading is strongly inhibited in the presence of RPA, whereas RPA depletion permits the accumulation of Rad51 together with multiple Rad51 paralogs. RPA is known to compete with Rad51 for ssDNA binding, thereby suppressing spontaneous nucleoprotein filament formation, and recombination mediators such as BRCA2 are required to promote Rad51 assembly on RPA-coated substrates (Jensen et al. 2010; Liu et al. 2010; Bhat and Cortez 2018). Our results are consistent with this framework and further suggest that RPA maintains ssDNA in a state that is refractory to premature recombination. The coordinated enrichment of Rad51 paralogs upon RPA depletion suggests that ssDNA becomes permissive for recombination rather than simply allowing nonspecific Rad51 binding.

Our findings also provide insight into the immediate consequences of RPA exhaustion during replication stress. Previous studies using human cells and purified proteins have shown that FBH1 interacts with PCNA, RPA, and Rad51, counteracts Rad51-mediated homologous recombination through the disruption of Rad51-filaments, and promotes DSB formation in response to replication stress (Fugger et al. 2009; Bacquin et al. 2013; Fugger et al. 2013; Jeong et al. 2013; Masuda-Ozawa et al. 2013; Simandlova et al. 2013; Tsutsui et al. 2014; Chu et al. 2015; Liu et al. 2023; Greer et al. 2026). Mechanistically, FBH1 catalyzes replication fork regression and facilitates recruitment of the MUS81 endonuclease, leading to cleavage of stalled replication forks (Fugger et al. 2013; Fugger et al. 2015). Interestingly, our data show that RPA depletion induces Fbh1 loading onto ssDNA, suggesting that RPA suppresses, rather than promotes, Fbh1 recruitment. It is therefore tempting to speculate that RPA exhaustion promotes FBH1 loading, which in turn facilitates fork regression and MUS81-dependent cleavage, ultimately leading to replication fork breakage. Notably, we did not detect obvious ssDNA fragmentation in RPA-depleted extracts, suggesting that additional steps or specific DNA structures may be required for fork breakage. Reconstitution of fork-like DNA structures in RPA-depleted NPE will be important for directly evaluating this model.

## Experimental procedures

### Preparation of *Xenopus* egg extracts

*Xenopus laevis* was purchased from Kato-S-Science and handled according to the animal experimental regulations at Kyushu University. NPE and HSS were prepared as described previously (Sparks and Walter 2018; Nishiguchi et al. 2025).

### Cloning and plasmids

Symbols of *Xenopus* genes and proteins conformed to the nomenclature guidelines of Xenbase (https://www.xenbase.org/entry/static/gene/geneNomenclature.jsp). The *Xenopus laevis rpa1* gene was amplified from *Xenopus* egg cDNA by PCR using primers 5′-GCAGGAACCATATGGCACTTCCACAACTGAGCG-3′ and 5′-GACGGAACAGATCTTCAGACACCTTGAGTTGCCA-3′, digested with NdeI (New England Biolabs, #R0111) and BglII (Takara Bio, #1021), and cloned into pDE1a (Kawasoe et al. 2016), resulting in pDE1a-xRPA1. The *Xenopus laevis rpa2* gene was amplified from *Xenopus* egg cDNA by PCR using primers 5′-GGAACTCGAGATGTGGAACAACCACGGTGGATTTG-3′ and 5′- GACAAGCTTTTAATCTCCATCAGTGCATTTGTAATGTTCATCATC -3′, digested with XhoI (New England Biolabs, #R0146) and HindIII-HF (New England Biolabs, #R3104), and cloned into pDE1a, resulting in pDE1a-xRPA2. The *Xenopus laevis rpa3* gene was amplified from *Xenopus* egg cDNA by PCR using primers 5′-GCAGGAACCATGGCGGATCTCTTCGATGTTCCTAAGG-3′ and 5′-GACGGAAGGATCCTCATTCGCTTGAACTGTGCCCAATGGG-3′, digested with NcoI-HF (New England Biolabs, #R3193) and BamHI-HF (New England Biolabs, #R3136), and cloned into pDE1a, resulting in pDE1a-xRPA3. Baculoviruses for the expression of Rpa1, Rpa2, and Rpa3 were constructed by transferring *rpa1*, *rpa2*, and *rpa3* genes into BaculoDirect C-Term Linear DNA (Life Technologies) using the Gateway LR reaction.

### Protein expression and purification

Purification of *Xenopus* RPA (Rpa1–3) was performed as follows: Recombinant proteins were expressed by co-infecting Sf9 insect cells with baculoviruses carrying *rpa1*, *rpa2*, and *rpa3* genes at 28°C in Sf-900 II SFM medium (Thermo Fisher Scientific, #10902104) supplemented with 2% (v/v) fetal bovine serum (FBS). Cells were harvested, washed with phosphate-buffered saline (PBS; 140 mM NaCl, 2.7 mM KCl, 10 mM PO_4_^3−^), and snap-frozen in liquid nitrogen. Cells were suspended in buffer H (25 mM Tris-HCl pH 7.4, 0.1 M NaCl, 1 mM dithiothreitol [DTT], 10% glycerol, 1 mM ethylenediaminetetraacetic acid [EDTA], 0.01% Nonidet P-40) containing 1 mM phenylmethylsulfonyl fluoride (PMSF), and 1× cOmplete, EDTA-free (Roche Life Science, #5056489001), and the lysates were centrifuged at 33,000 rpm for 30 min in a Beckman 70Ti rotor. Cleared lysates were loaded on a HiTrap Q HP column (Cytiva, #17115301), and bound proteins were eluted with a 0.1–1 M NaCl linear gradient with buffer H. Peak fractions (2 mL) were pooled and diluted with 5 mL of buffer H containing 0.3 M NaCl, mixed with ssDNA agarose beads, rotated for 90 min at 4°C, and the bound proteins were eluted with buffer H2 (25 mM Tris-HCl pH 7.4, 1.5 M NaCl, 1 mM DTT, 10% glycerol, 1 mM EDTA, 0.01% Nonidet P-40, 50% ethylene glycol). The peak fractions were pooled, passed through a PD-10 desalting column (Cytiva, #17085101) equilibrated with buffer H, loaded on a Mono Q 5/50 GL column (Cytiva, #17516601), and bound proteins were eluted with a 0.1–1 M NaCl linear gradient with buffer H. Peak fractions were applied to a Superdex 200 Increase 10/300 GL (Cytiva, #28990944) column equilibrated with buffer H lacking Nonidet P-40, and the eluted fractions were snap-frozen in small aliquots in liquid nitrogen and stored at −80°C.

### Immunological methods

Production of the antibody against *Xenopus* Msh6 was described previously (Kawasoe et al. 2016). Rabbit antisera against *Xenopus* RPA and Rad51 were raised against the untagged Rpa1–3 complex expressed in Sf9 cells and N-terminally His_6_-tagged Rad51 expressed in *E. coli*, respectively. The expression and purification of *Xenopus laevis* Rad51 are described elsewhere. For immunoblotting, the antisera were used at a dilution of 1:10,000.

For immunodepletion from NPE, 3 vol of RPA antisera were bound to 1 vol of recombinant Protein A Sepharose Fast Flow (PAS; Cytiva) overnight at 4°C. Antibody-coupled PAS beads (0.2 vol) were then incubated with 1 vol of NPE at 4°C for 1□h, and this procedure was repeated twice. Typically, 20–40□μL of extract was used per experiment.

### Preparation of primed circular ssDNA

Circular phagemid ssDNA was prepared using the M13KO7 helper phage as described previously (Higashi et al. 2012). A 20-mer synthetic oligonucleotide (5′-TCGGAGGACCGAAGGAGCTA-3′) was annealed to pBluescript II KS(−) ssDNA in Tris-EDTA buffer (TE buffer: 10 mM Tris-HCl pH 7.4, 1 mM EDTA) containing 0.1 M NaCl.

### DNA synthesis in *Xenopus* egg extracts

HSS and NPE were supplemented with 2□mM adenosine triphosphate (ATP), 20□mM phosphocreatine (PC), and 5□μg/mL creatine phosphokinase (CPK), and HSS was further supplemented with 3□μg/mL nocodazole. The extracts were pre-incubated at 22°C for 5 min, followed by the addition of 0.25 vol of DNA substrates. DNA concentrations are indicated in the figures and legends. To monitor DNA synthesis, NPE was supplemented with 25 μM Cy5-conjugated deoxyuridine triphosphate (Cy5-dUTP; Cytiva, #PA55022) together with 50 ng/μL aphidicolin (Sigma-Aldrich, #A0781) or dimethyl sulfoxide (DMSO). The samples were incubated at 22°C for 2 h, and aliquots (3.0–3.6□μL for most experiments) were sampled and stopped with 100□μL of 1% sodium dodecyl sulfate (SDS) in 20□mM EDTA. DNA was purified by treatment with 50□μg/mL proteinase K, extracted with phenol/chloroform, precipitated with ethanol, and dissolved in TE buffer containing 10□μg/mL RNaseA at 2 ng/μL. DNA was separated on a 0.8% agarose gel, followed by staining with SYBR Gold nucleic acid gel stain (Thermo Fisher Scientific, #S11494). Fluorescent signals were acquired using the Amersham Typhoon Scanner 5 system (Cytiva) and processed using Fiji (https://fiji.sc).

### The ssDNA pulldown assay

A 90-mer synthetic oligonucleotide (5′-AAATAGACAGATCGCTGAGATAGGTGCCTCACTGATTAAGCATTGGTAACTGTCAGACCAAGTTTACTCATATATACTTTAGATTGATTT-3′; Eurofins Genomics), conjugated with a biotin-ON moiety at the 5′ terminus, was mixed with streptavidin (SA; New England Biolabs, #N7021S) at final concentrations of 50 nM DNA and 50 μg/mL protein in buffer W (10 mM Tris-HCl pH 7.4, 1 mM EDTA, 1 M NaCl, and 0.1% Triton X-100) at 4°C with gentle rotation to form DNA–SA complexes. The DNA–SA complexes were then immobilized on biotinylated Sepharose beads at 0.25 pmol DNA per μL beads at 4°C with gentle rotation in buffer W (Higashi et al. 2012). The beads were washed with buffer W twice, and then with ELB salts three times (100 mM HEPES-KOH pH 7.8, 0.5 M KCl, and 25 mM MgCl_2_). The beads were added to NPE at 0.33 μL beads per μL extract, and the mixture was incubated at 22°C for 10 min with gentle rotation. After incubation, the beads were washed with ELB salts three times, and bound proteins were extracted with 1×SDS sample buffer (50 mM Tris-HCl pH 6.8, 0.1 M dithiothreitol, 2% SDS, 0.05% bromophenol blue, and 10% glycerol). Typically, 20–40□μL of extract was used per experiment.

### LC-MS/MS analysis

Protein samples were separated on a 4–20% Mini-PROTEAN TGX precast gel (Bio-Rad, #4561096) and stained with GelCode Blue Stain Reagent (Thermo Fisher, #24590). The gel region containing proteins of interest was excised and cut into approximately 1-mm pieces, and proteins in the gel pieces were reduced with 10 mM DTT (Thermo Fisher Scientific, A39255) in 25 mM ammonium bicarbonate (FUJIFILM Wako, 018-21742), alkylated with 55 mM iodoacetamide (Thermo Fisher Scientific, A39271) in 25 mM ammonium bicarbonate, and digested overnight at 37°C with trypsin and lysyl endopeptidase (Promega, V5073) in a buffer containing 40 mM ammonium bicarbonate pH 8.0. The digested peptides were extracted from gel pieces with 50% acetonitrile (FUJIFILM Wako, 012-19851) containing 0.1% formic acid (FUJIFILM Wako, 067-04531), followed by 70% acetonitrile containing 0.1% formic acid, combined in a fresh vial, and concentrated to 15 μL in a centrifugal evaporator. The resulting peptides were analyzed on an Advance UHPLC system (AMR/Michrom Bioresources Inc) coupled to a Q Exactive mass spectrometer (Thermo Fisher Scientific), and the raw mass spectra were processed using Xcalibur 4.0.27.19 (Thermo Fisher Scientific). Peptides were separated on a micro-ODS column (L-column 2 ODS, 3 µm, 0.1 x 150 mm, PEEK-steeved type, CERI) at 40°C and a flow rate of 500 nL/min using a 90-min gradient in which solvent B (acetonitrile, 0.1% formic acid; FUJIFILM Wako) was increased from 5% to 35%. Raw LC-MS/MS data were searched against the UniprotKB protein database restricted to *Xenopus laevis* using Proteome Discoverer version 1.4 (Thermo Fisher Scientific) with the Mascot search engine version 2.7 (Matrix Science). A decoy database comprising either randomized or reversed sequences from the target database was used to estimate the false discovery rate (FDR), and the Percolator algorithm was used to evaluate false positives. Search results were filtered to a global FDR of 1% to obtain high-confidence protein identifications.

### Mass spectrometry data analysis

Total spectrum count values were extracted from Scaffold version 5 (Proteome Software). Proteins detected in at least 3 out of 6 samples (2 conditions with 3 biological replicates each) were subjected to further analysis. The data were log_2_-transformed prior to statistical analysis, and missing values were imputed by random draws from a Gaussian distribution N(μ − 1.5σ, (0.3σ)²), where μ and σ are the mean and standard deviation of the observed log_2_-transformed values, under the assumption that missing values predominantly arise from low-abundance proteins (Tyanova et al. 2016). Log_2_ fold changes were calculated between conditions, and statistical significance was assessed using Welch’s t-test. P values were converted to −log_10_ values for visualization.

## Data availability

All data from this study are included in the manuscript and supporting information. Unprocessed raw mass spectrometry data are available at https://doi.org/10.48708/7430646, and other raw data will be shared on reasonable request to the corresponding author.

## Acknowledgments

We thank Toshiki Tsurimoto for critical reading, Shota Fukuda for assistance with the preparation of recombinant Rad51 protein and antisera, and the members of the Takahashi laboratory for helpful discussions. This work was supported by the program of the Joint Usage/Research Center for Developmental Medicine and High Depth Omics, IMEG, Kumamoto University.

## Funding and additional information

This work was supported by MEXT KAKENHI [grant number 24H02328], JSPS KAKENHI [grant numbers 24K01956, 22H04697, 20H05392, 20H03186 to T.S.T., 24K09328, 22K15036 to Y.K.], and JST SPRING [grant number JPMJSP2136 to T.M.].

## Author contributions

T. M.: Conceptualization, Formal analysis, Investigation, Methodology, Validation, Visualization, Funding acquisition, Writing—original draft.

N. T.: Investigation, Resources

Y. K.: Methodology, Funding acquisition, Writing—review & editing.

K. I.: Investigation, Resources

T. S. T.: Conceptualization, Funding acquisition, Supervision, Methodology, Writing—original draft, Writing—review & editing.

## Abbreviations

ATM: Ataxia-telangiectasia mutated
ATR: Ataxia telangiectasia and Rad3-related protein
Cy5-dUTP: Cy5-conjugated deoxyuridine triphosphate
ETAA1: Ewing tumor-associated antigen 1
Fbh1: F-box DNA helicase 1
HSS: high-speed supernatant
NPE: nucleoplasmic extracts
RF-A: Replication factor A
Rfwd3: RING finger and WD repeat domain 3
RPA: Replication protein A
ssDNA: single-stranded DNA
SA: Streptavidin
xRPA: *Xenopus* RPA
Xrcc3: X-ray repair cross complementing 3.

---

**Supplementary Table S1. Proteins identified by mass spectrometry in the ssDNA pull-down assay**

Summary of the mass spectrometry analysis. Total spectral counts for the indicated proteins are listed together with their molecular masses.

